# Modelling sterilisation strategies to maximise population impact and cost-efficiency in free-roaming dog populations

**DOI:** 10.64898/2026.07.02.735983

**Authors:** H.R. Fielding, P.R. Bessell, A.D. Gibson, K. A. Fernandes, V.R. Amulya, L. Gamble, B.M.de C. Bronsvoort, I.G. Handel, R. King, R.J. Mellanby, S. Mazeri

## Abstract

Free-roaming dog populations pose challenges to public health and animal welfare, and their management and protection is legally mandated in many countries to control zoonotic diseases such as rabies, reduce bites, mitigate fear of aggressive dogs, and preserve animal welfare. Although international organisations recommend holistic dog population management with surgical sterilisation as a key component, stakeholders report a lack of evidence-based guidance on effective and cost-efficient strategies.

To address this, we developed a deterministic model of a closed free-roaming dog (FRD) population, including dogs dependent on and independent of humans, parameterised using data from southern India. We evaluated a wide range of sterilisation strategies varying duration, interval, frequency, intensity, targeting different population subsets, and fixed or responsive approaches. Strategies were assessed based on their effects on independent FRD population size and implementation costs.

Female sterilisation coverage (FSC) strongly determined population reduction and cost-effectiveness. Repeated, short sterilisation periods with appropriately timed intervals and high early intensity best balanced population reduction and cost-efficiency. Excluding dependent dogs (cared for by humans) severely constrained population reduction, while male sterilisation enhanced impact at little extra cost, assuming direct effects on pregnancy rates. Costs and population impacts can be visualised on an interactive web application (https://field.shinyapps.io/DogPopSimApp), allowing practitioners and policymakers to explore sterilisation strategy outcomes. Overall, cost-effective population reduction required sustained commitment and coverage-driven strategies that balance welfare, population reduction and the operational needs of dog-mediated rabies control.

**Author Summary:** Free-roaming street dogs live closely with people. While these animals are often valued members of the community, they can raise public health and safety concerns, including disease transmission, bites, and traffic accidents. To protect both human and animal health, many regions use surgical sterilisation to humanely manage street dog populations. However, running these programmes is complex and expensive, and community managers often lack clear guidance on how to maximise their limited resources.

To address this, we used data from southern India to create a mathematical model of a street dog population. We simulated thousands of different sterilisation strategies to see how factors like campaign length, frequency, and targeted sub-groups impacted population size and total costs.

We found that short, repeated sterilisation campaigns offer the best balance of population reduction and cost-efficiency. Crucially, these programmes must include dogs that are partially cared for by humans; otherwise, their puppies will continually replenish the street population. Although sterilisation of females drives population reduction, sterilising males alongside females enhanced the overall impact. Ultimately, our study provides practical, evidence-based guidance to help communities design effective and financially sustainable dog management programmes that improve public health and animal welfare.

## Introduction

In many countries, domestic dogs freely roam and live alongside human populations as part of society [1]. However, this can cause conflict, particularly in countries with high free-roaming dog populations, such as India. In 2023, there were 2.76 million reported dog bites in India [2]. In rabies-endemic countries such as India, bites are not only damaging in themselves but carry the extra risk of rabies transmission. Domestic dogs can be a source of several major zoonotic diseases (including rabies, leishmaniasis, echinococcosis, toxocariasis), and free-roaming domestic dogs (FRDs) may depredate livestock, transmit disease and threaten wild species where habitats overlap. In regions with large FRD populations and where conflicts are common, there is often a desire to reduce the FRD population and mitigate these negative impacts.

Populations can be reduced by restricting food resources, removing animals (via culling, sheltering or adoption) or restricting fertility. Evidence suggests that although culling initially reduces population size, it is ineffective as a long-term solution due to compensatory ecological mechanisms, may initially increase disease transmission, and many societies find it unacceptable [3,4]. As a sole intervention, restricting food resources is likely to compromise welfare in the short-term and is complex in a society where feeding free-roaming dogs is common. Though combined with other measures to reduce the population size, it may be effective in reducing carrying capacity and limiting population rebound via increased fecundity or immigration. Alongside responsible pet ownership, surgical sterilisation is recommended by the Animal Welfare Board of India and at the time of writing has been notified by the Government of India and mandated by the Supreme Court of India as the preferred method to control India’s FRD population [5,6]. Sterilisation is suggested to stabilise and reduce the population and maximise the efficiency and impact of rabies vaccination campaigns [7] and may reduce dog bites via reduced dog numbers and/or reduced reproductive behaviours [8]. If sterilisation is performed in an effective and humane manner, it is likely to improve FRD welfare and longevity via increased resources and reduced energy spent on reproduction [9,10]. Surgical sterilisation has been associated with increased body condition and better health in FRDs [11,12] and may reduce contact rate through reproductive-related behaviours, as identified in possums [13]. However, surgical sterilisation is labour intensive, costly and may compromise welfare if guidelines are not adhered to. In addition, the behavioural impacts of sterilisation on FRDs are unclear. Sterilisation is often inconsistently applied over short periods or at low levels, which is unlikely to reduce population size [14]. When moderate intensity sterilisation (58-66% sterilisation coverage) was applied in a single pulse, empirical data showed the population stability increases and individual animal welfare improved, but no impact on population size was seen, at least over a two-year follow-up period [11]. Modelling in free-roaming cat populations showed that low intensity sterilisation (over time achieving sterilisation coverage of ∼60%) was less likely to achieve population reduction and did not minimise preventable deaths compared to high intensity sterilisation [9]. Modelling and empirical studies have demonstrated the intense sterilisation effort necessary to effect population change [15,16], yet via a user involvement approach, Collinson et al. found that international stakeholders with experience working in dog population management lacked evidence-based recommendations for how to implement sterilisation campaigns that reduce the size of the dog population [17].

To assist stakeholders in planning sterilisation campaigns we used field data to construct a deterministic mathematical model of free-roaming dogs representative of urban Southern India. We applied 1524 surgical sterilisation strategies to the simulated population, altering duration of sterilisation, intervals between sterilisation (both fixed intervals and in response to number of dogs captured ‘catch-guided’), time of year, and targeting different subsets of FRDs. We assessed their success based on population density change and cost of campaign, puppy births and deaths, adult FRD deaths, vaccinations required for rabies control campaigns and population turnover. We recorded the total number of sterilisations and sterilisation coverage to better understand what values were associated with optimal strategies. All strategies applied were logistically feasible for the population size that the model is based on, based on authors experience. This paper aims to explore which features of sterilisation strategies reduce population size in a cost-effective manner. Particular areas of interest to end users outlined by Collinson et al. were to explore; the proportion of the population that needs to be sterilised and over what period, the effects of sterilisation strategies of different size and duration, the effects of single sterilisation strategies, the optimal frequency for repeated strategies and the effect of female-only strategies. We explicitly included seasonal variation of reproduction, previously identified in FRDs [18], to see how this may affect the success of a sterilisation strategy, which has also been identified as of interest in dog population management [19].

## Methods

### Baseline model

We constructed a deterministic mathematical model simulating a FRD population comprised of dependent (have provisions of shelter, food, and/or care by humans) and independent dogs (spend all their time unconfined, no person considers themselves an owner of these dogs, yet varying levels of food may be provided by humans) that have access to public spaces (Fig. 1). Confined dogs (dogs that are never allowed to roam freely) are not included in this model. The model population size and structure (age, % female, % dependent; Table S1) is based on data from Margao municipality (Goa, India) collected during Mission Rabies vaccination campaigns from 2015 to 2021, [20]. Margao has an estimated area of 16.37km^2^ and an estimated human population of 113,000 based on 2021 predictions from 2011 census data (2021 data unavailable due to COVID-19 pandemic) [21], giving an estimated human:dog ratio of 17.25. This municipality is typical of urban Goa in terms of FRD demography. FRD population estimates showed little variation over five vaccination campaigns (2015-2021; mean = 6553, SD = 504) therefore we assumed the population, which had minimal dog population management during these years, was stable around its carrying capacity, and fitted the model to replicate this. The starting model population size was 6500 for all simulations. Overall carrying capacity was therefore 6500 and for respective sub-groups (e.g. dependent dogs) the carrying capacity was set as the starting population size for that sub-group. Based on field data, the dependent FRD population comprised one third of the total model population and was split by a sex ratio of 2.5 males to 1 female. The rest of the population was independent and split by a sex ratio of 1.5 males to 1 female, reflecting differences in the sub-group structure. The total model population is closed without immigration or emigration, as we did not have accurate estimates to suggest the direction of net movement. We acknowledge that the free-roaming dog population does not discretely fit into these two populations and that there are many dogs that lie on a spectrum between the two or that switch groups with adoption or abandonment, however for modelling purposes and based on the classification of our data, we felt this was a reasonable grouping. Juveniles and puppies could move from the dependent to the independent population through abandonment (Fig. 1) and the proportion of sterilised males from the independent and dependent populations affected pregnancy rates in both sub-populations, therefore breeding effectively spanned both populations.

**Figure 1.**
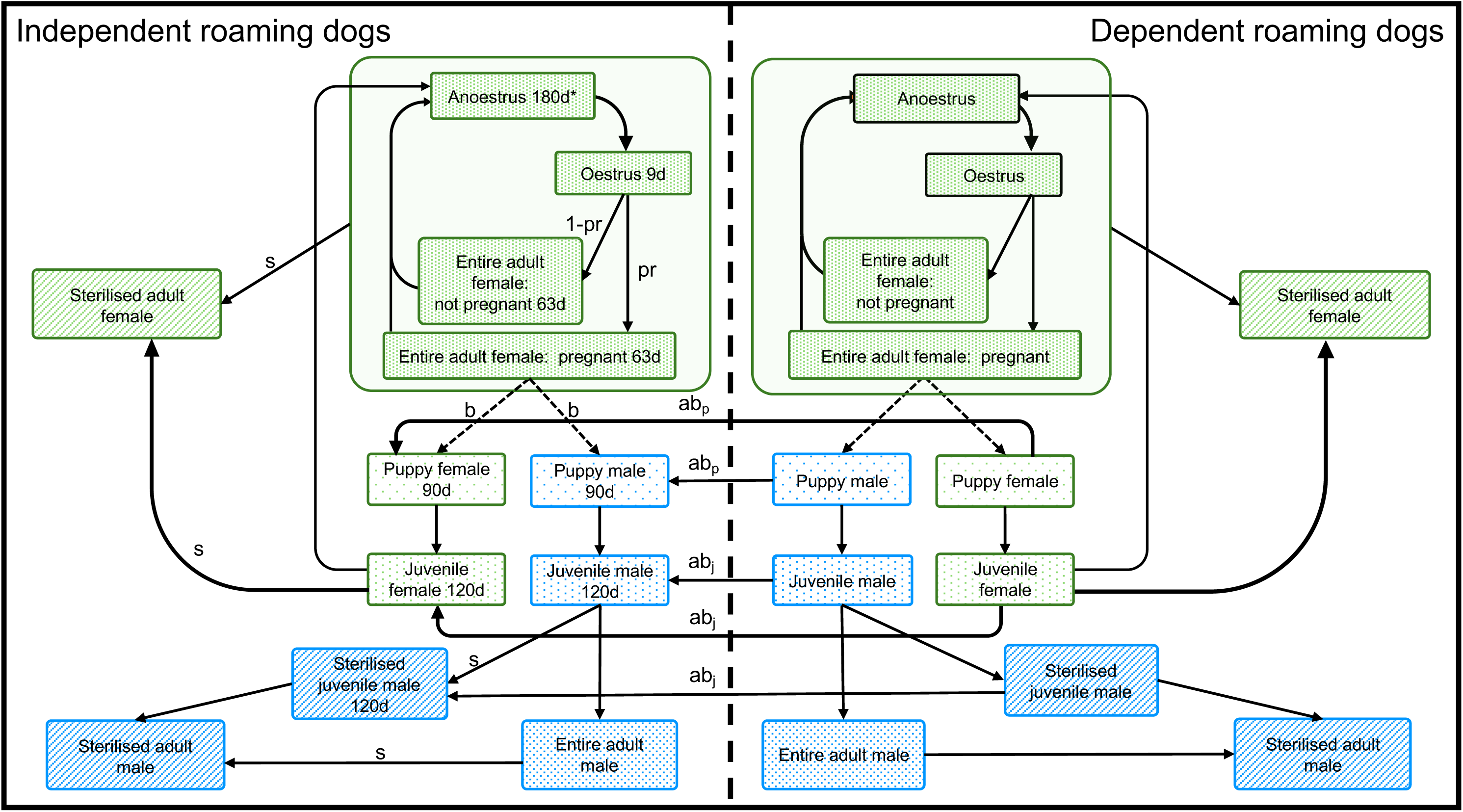
Model schematic. Schematic representation of the life cycle of free-roaming dogs (FRDs) in the simulated dependent and independent populations in an urban Indian setting. The deterministic model divides dogs by dependency status, sex, age and reproductive or sterilisation status. (s), sterilisation rate during intervention periods, determined by catching effort and the proportion of dogs already sterilised; (pr), probability that a female becomes pregnant following oestrus (altered by a factor relative to the total proportion of sterilised males); (b), six puppies produced per pregnancy, with a 1:1 male-to-female ratio; (ab_p_) and (ab_j_), abandonment rates through which dependent puppies and juveniles, respectively, enter the independent population. Compartments with limited duration have the number of days spent in that state listed in days. *Seasonal breeding is represented by a 2.5-fold increase in the rate at which females leave anoestrus between days 225 and 315 of each year. Deaths occur from all compartments according to baseline mortality rates specific to age, sex and dependency status. In the independent population, mortality is additionally adjusted according to the current independent population size relative to its initial size (assumed carrying capacity). Rates and timings apply to both population sub-groups but are only labelled in the independent population to improve visualisation.

### Mortality and abandonment

Mortality and abandonment parameters were fitted to achieve a stable population at carrying capacity (*K)* that had a similar sex and age structure to our empirical data (Table S1) without interventions applied (the baseline model). Adults, juveniles, puppies, female and male, dependent and independent dogs have different mortality rate based on life expectancy estimates of female dependent dogs [22] assuming that dependent and male dogs live longer than independents and females respectively. There is no change in mortality after sterilisation. In the independent population only, mortality rates were adjusted according to:

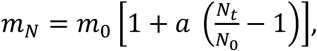

where *m_N_* is the adjusted mortality rate, *m_0_* is the baseline mortality rate when *N_t_* = *N_0_*, *N_0_* is the initial independent population size, *N_t_* is the current independent population size, and *a* is the adjustment factor (0.5). With *m*_0_ = 0.1 and *N_0_* = 5000, *m_N_* would be increased to 0.15 at *N_t_* = 10,000 but decreased to 0.075 at *N_t_* = 2500. Whilst there is likely to be some adoption from the independent to dependent population, we assume the net movement between puppy compartments was abandonment from dependent to independent populations. The abandonment rate from the dependent to the independent puppy and juvenile population was adjusted according to population size relative to reference population thresholds. For puppies, abandonment was proportional to the ratio of the current dependent puppy population to a puppy ceiling, defined as 12% of the initial dependent population. For juveniles, abandonment was proportional to the ratio of the current dependent population to the initial dependent population. Thus, abandonment decreased when the relevant population was below its reference threshold and increased when it exceeded it. Mortality and abandonment values were similar to those used in previous modelling [23].

### Age and reproduction parameters

Dogs are considered puppies from 0−90 days, juveniles from 91−300 days and adults over 300 days. Female dogs enter their first oestrus as they become adults at 300 days (Table S1), they are in oestrus for 9 days and then go into a 63-day period of pregnancy (35%) or pseudo-pregnancy (65%), after which they enter anoestrus for 180 days (Fig. 1). Pregnant females birth 6 puppies with 50:50 sex ratio (Table S1). During a period of the year which is defined as ‘season’, the rate of dogs coming out of anoestrus is accelerated by a factor of 2.5, to simulate the seasonal pregnancy prevalence data from independent and dependent dogs submitted for sterilisation at the Worldwide Veterinary Service (WVS) clinic in North Goa (Fig. S1), where breeding was observed to increase between mid-August to mid-November [18].

### Sterilisation

Dogs were eligible for sterilisation as juveniles or adults. To construct realistic sterilisation strategies, the authors utilised recent experience in operating fixed and mobile sterilisation clinics in Goa to estimate realistic capacity of catching and sterilisation. The estimated maximum number of dogs that could be caught (and therefore sterilised) in one day in this simulated area was 125 (5 catching teams each catching 25 dogs per team during 2 sessions in one day) [24]. As sterilisation coverage increased, fewer unsterilised dogs were captured, therefore, sterilisation rate was constrained by increasing sterilisation coverage in each respective subpopulation. There was no distinction between sterilisation and catching; if a dog was caught, it was sterilised. Catching was performed 5 days per week. We allowed some strategies to respectively decrease (no less than 1) or increase (up to 5) the number of catching teams if on average teams caught fewer than 10 dogs or more than 90% of the maximum possible dogs on the previous day. This followed the typical strategy used in the field to minimise staff and transport costs. We conservatively assumed 5% of independent dogs were uncatchable [15] and 10% of dependent dogs would not be sterilised due to guardian choice. Sterilisation started after a warm-up period of six years and the simulation ran for a subsequent 10 years.

Six main types of strategy were tested; 1 – ‘catch-guided’, where sterilisation was stopped for fixed intervals (30, 90, 150, 180, 365, 730, 1825 days) if the total number of captured dogs (i.e. not per team) was below a threshold (5 or 10 dogs), 2 – ‘fixed-maintain’ strategies with pre-defined intervals (90, 185, 365, 730, 1825, 2555 days), duration (30, 90, 180, 365, 730, 1825, 2555 days) and repeat sterilisation periods (1, 3, 5 or repeated until end of strategy), 3 – ‘fixed-reduce’ same as ‘fixed’ but catching teams are reduced if the mean number of dogs captured per team is less than 5 or 10, 4 – ‘single-session’ a single sterilisation period of fixed duration (30, 90, 180, 365, 730, 1825, 2555, 3650 days), 5– ‘continuous’ sterilisation continued throughout the 10 year study period, and 6 – ‘seasonal’ annual campaigns of 6-months-on 6-months-off varying the time of year that sterilisations started. All these strategies were tested with all combinations of female-only or male-inclusive sterilisation, independent-only or dependent-inclusive sterilisation and with 1, 3, or 5 initial catching teams. FSC was calculated at every timestep (1 day). To summarise the entire simulation, the mean-daily FSC was calculated by summing each daily coverage and dividing by the number of days in the model.

Given that domestic dogs exhibit mate selection, where unfamiliar males are less successful and females reject some males [25–27], male sterilisation is likely to reduce pregnancy rates to some extent, though empirical evidence is lacking. Due to uncertainty around this assumption, we tested this parameter (male sterilisation impact) with values of 0, 0.15 and 0.25 for all strategies. For the main analysis, we selected 0.15 as a reasonable estimate male sterilisation impact, where 70% male sterilisation resulted in a decrease in pregnancy rates from 10 to 9%.

### Costs

Costs were taken from a WVS sterilisation campaign in Goa in 2022 in a pop-up clinic only used for the purpose of sterilisation, i.e. all overheads are charged to the campaign (Table S2) and were converted from Indian rupees to US dollars using an exchange rate of 0.014 from 2022. In the authors’ experience, an experienced surgical vet can perform up to 15 sterilisations today, reducing to 10 if only females are caught due to the increased surgical time for ovariohysterectomy (Table S2). Reported costs are for comparison purposes within this study and may not translate to other contexts.

### Outcome measures

Strategies were categorised depending on their cost and effect on the population and both were discounted annually at the recommended rate of 3%[28]. Population density and puppy births and deaths, and adult deaths were highly positively correlated (∼0.9) among strategies (Fig. S2), therefore we use population density as a proxy for all population measures. The cost threshold was set at United States $250,000, equal to a per person annual spend of $0.22 over 10 years (based on Margao population size), strategies above this were categorised as ‘high-cost’ and below as ‘low-cost’. The population impact was deemed ‘effective’ if the final number of independent dogs over 3 months old in year ten of the simulation was reduced by 50% or more and the population density was decreased by more than 40% over the 10-year simulation.

For each input variable value (e.g. one input variable was number of teams at the start and the values tested were 1, 3 and 5 teams), we calculated the proportion of strategies that were optimal and conducted pairwise comparisons using Fisher’s exact test with an ‘fdr’ correction (Benjamini and Hochberg, 1995) to evaluate differences between the proportions using the ‘fisher.multcomp’ function from the R package ‘RVAideMemoire’ [30]. Modelling, data wrangling and visualisation was performed using R [31] and the ‘tidyverse’ [32].

## Results

Out of 1524 tested strategies, forty-one were categorised as low-cost and effective and henceforth referred to as ‘optimal’ (Fig. 2). Summaries of cost and population density reduction for all 1524 strategies are outlined in Table 1 and example strategies are shown in Figure 3.

**Figure 2.**
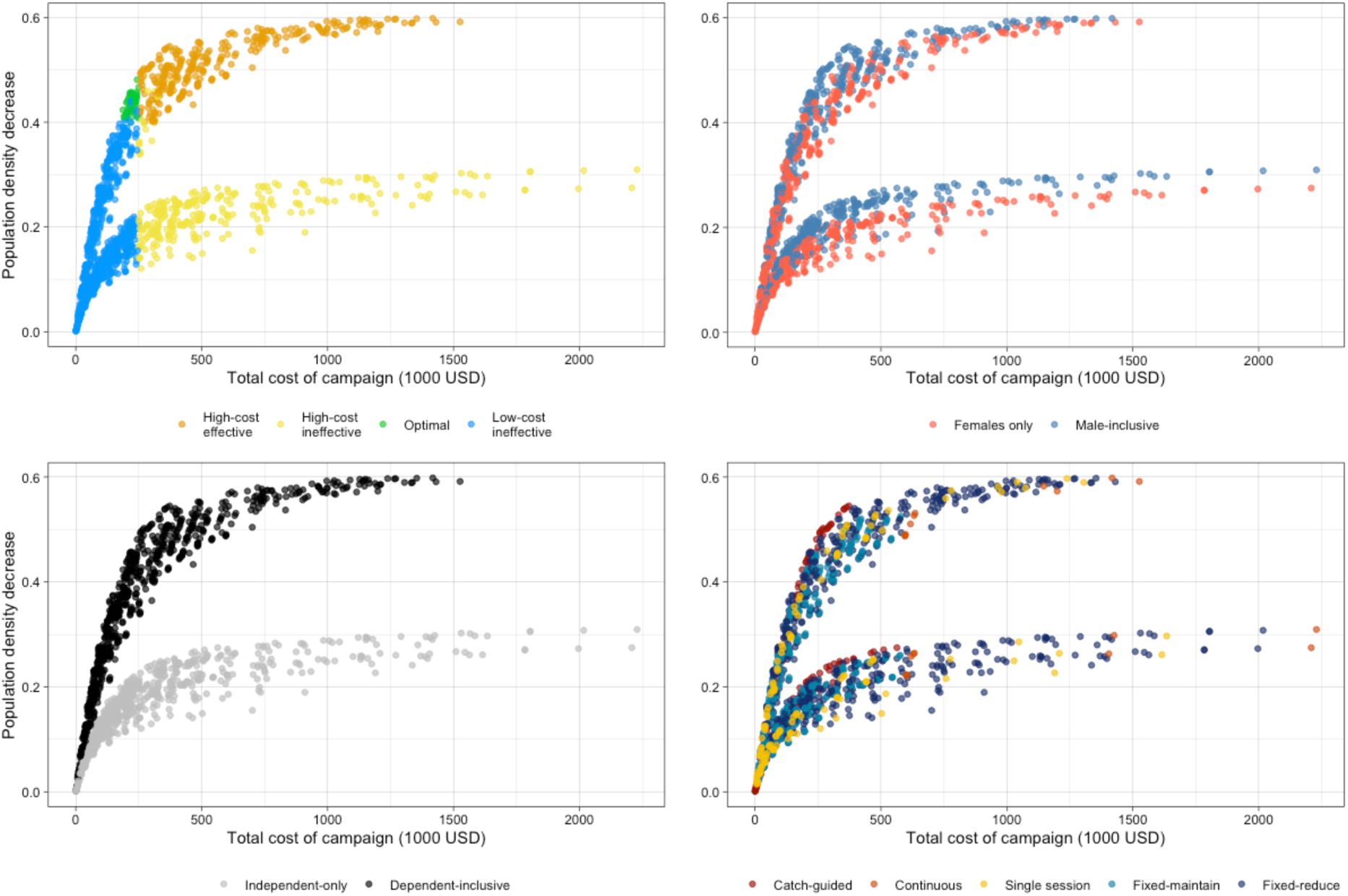
Strategies overview. Free-roaming dog (FRD) total population density change and cost of sterilisation strategies over 10 years in the simulated FRD population. Top left points coloured by strategy success based on cost (high > US$250k, low <= 250k) and population density reduction (effective = more than 40% density reduction and less than 50% population at 10 years) with low-cost effective strategies classed as optimal, top rightpoints coloured by female-only and male-inclusive strategies, bottom left by independent-only and dependent-inclusive (including FRDs cared for by humans) FRDs and bottom right panel by type of strategy approach. Seasonal strategies are not included as they represented the same strategy, only varying by month of start.

**Figure 3.**
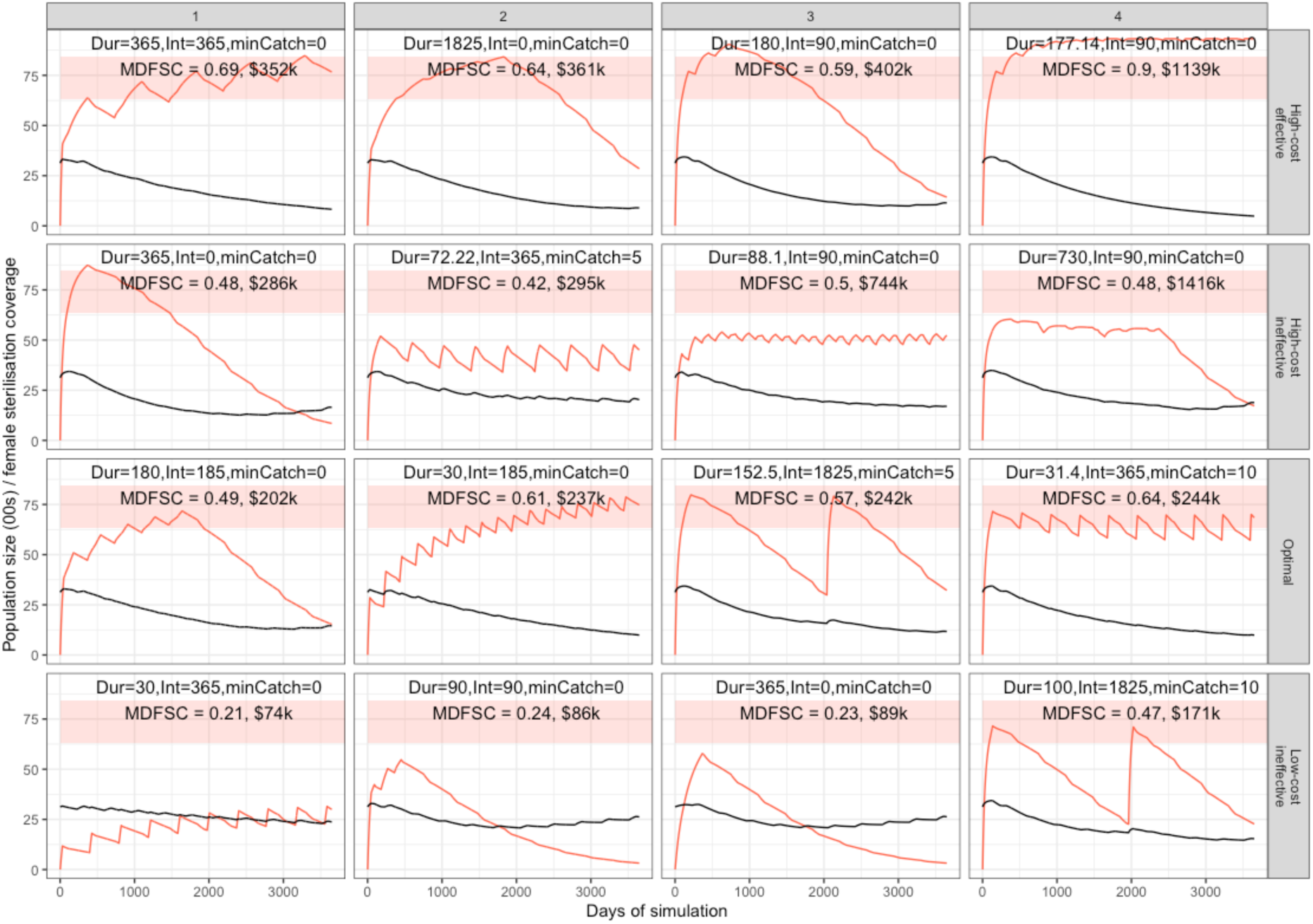
Example strategy outcomes. Female sterilisation coverage (FSC; red line) and free-roaming dog population size (black line) over ten years for four sterilisation strategies in each success category based on cost (high > US$250k, low <= 250k) and population density reduction (effective = more than 40% density reduction and ineffective = less than 50% population at 10 years). Low-cost and effective strategies classed as ‘optimal’. The optimal zone for maximum FSC is indicated by the red shaded region. Individual plot annotations show the length of sterilisation in days (Dur, mean duration if catch-guided strategy), interval between sterilisation periods in days (Int), threshold number of dogs to stop catching (0 if fixed sterilisation duration), mean-daily FSC (MDFSC) and the total cost of the strategy. Plots are arranged from lowest to highest cost from left to right in each success category.

**Table 1.**
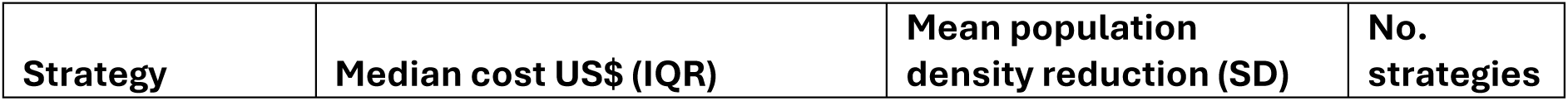
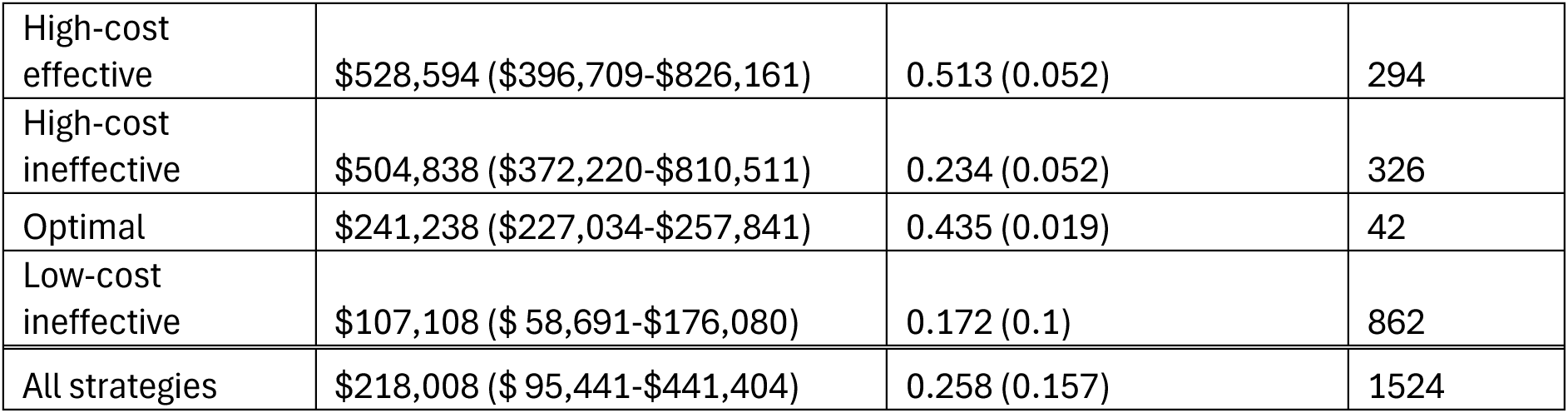
Summary statistics for strategies in each strategy group and overall for cost and population density reduction.

### Sterilisation coverage

Optimal strategies had mean-daily FSC between 47.5% and 64.3% (mean = 55.8%, SD = 4.56%) and a maximum FSC of between 63.3 and 84.5%. (mean = 74.1%, SD = 6.10%; Fig. 4). Mean-daily FSC and max FSC were highly correlated with each other (0.907, 95% CI = 0.898-0.916). Some optimal strategies had a rising population from the last lowest population size during the simulation (Fig. S9) and were closer to the population reduction cut-off of 40%, suggesting these strategies may not have been effective given a longer study period. These were all male-inclusive strategies and on average were associated with lower mean-daily FSC (population rising = 52.6%, SD = 3.0%, population static/decreasing = 59.7%, SD = 2.7%). Strategies with a max FSC below 63.3% were all ineffective. Optimal strategies spent a mean of 32.7% of days over the lowest maximum FSC (63.3%), whereas high-cost effective strategies spent a mean of 79.4% of days over this value. Of the strategies with a max FSC above 63.3%, high-cost ineffective strategies tended to achieve maximum FSC towards the start or end of the simulation, suggesting they did many sterilisations early on but then did not perform sufficient sterilisations to maintain the sterilisation coverage or did not achieve high FSC early enough to make sufficient impact in the population. Increased mean-daily FSC was highly correlated with population decrease (Spearman’s ρ = 0.953, 95% confidence intervals (CI) = 0.948-0.957), but less strongly correlated with costs (Spearman’s ρ = 0.677, 95% CI = 0.649 – 0.703). There was a strong negative relationship between number of FRD deaths and mean-daily FSC, even at low sterilisation coverage (Fig. S3).

**Figure 4.**
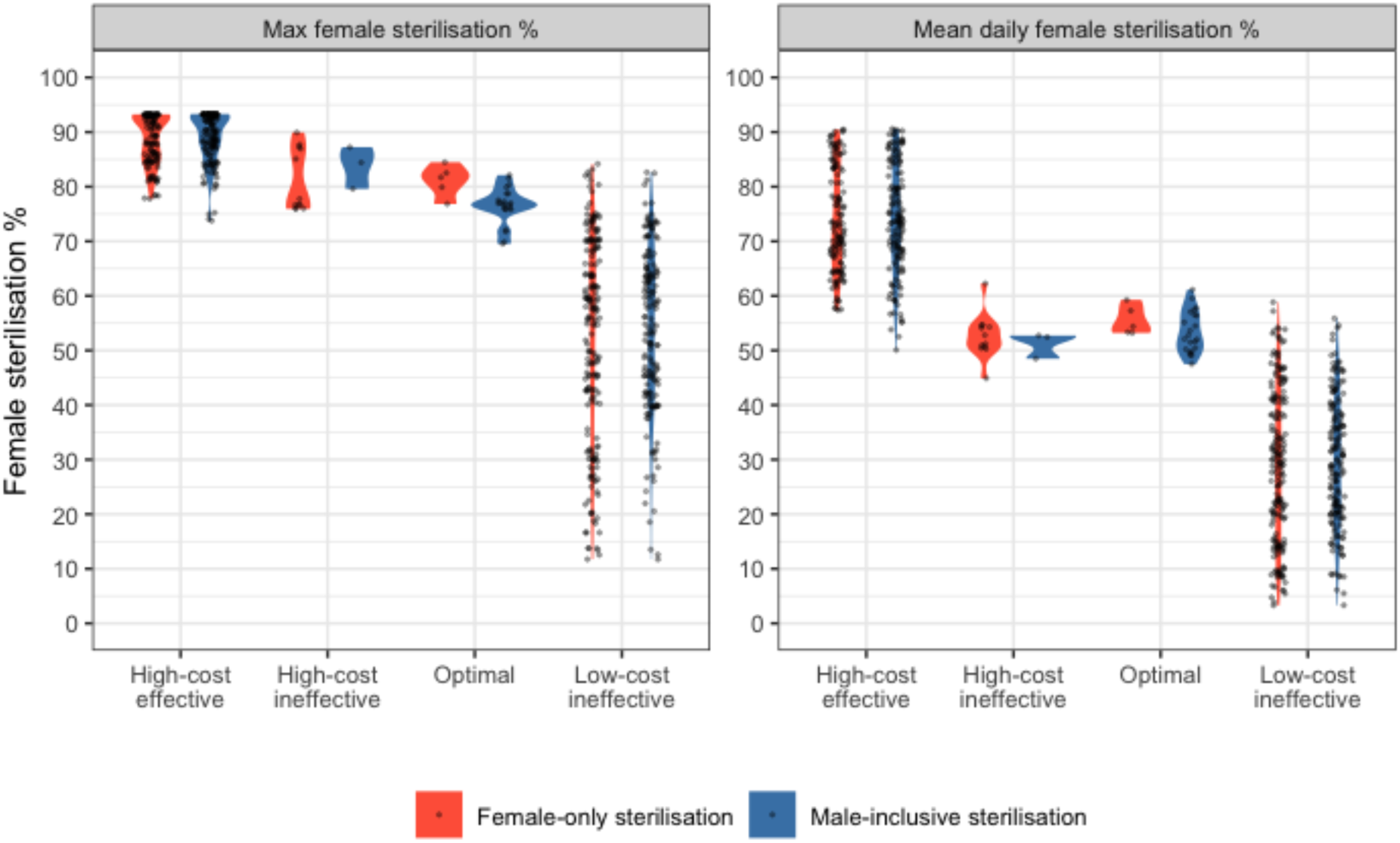
Sterilisation coverage and strategy success. Maximum (left) and mean (right) daily female free-roaming dog (FRD) sterilisation percentage (FSC) achieved in the 10-year simulation for each of 1524 strategies. Colour of background violin plot indicates whether the strategy sterilised males and strategies are split by their success based on cost (high > US$250k, low <= 250k) and population density reduction (effective = more than 40% density reduction and less than 50% population at 10 years).

### Sterilising dependent FRD

When only the independent population was sterilised, population reduction was limited and all strategies were ineffective, yet in dependent-inclusive strategies, 5.99% were optimal (Fig. 2, Table S3). Some independent-only strategies achieved a relatively high maximum FSC of 60.7% (mean max FSC = 44.6%, SD = 11.9%) as catching capacity was focussed on independent dogs, though still lower than max FSC achieved in dependent-inclusive strategies (mean max FSC = 67.3, SD = 22.5%). However, not being able to sterilise dependent FRDs, which made up one third of the FRD population, meant the mean-daily FSC was not sufficient to sustainably curb the population (independent-only mean-daily FSC mean = 29.9%, SD = 13.2% c.f. dependent-inclusive mean = 48.7%, SD = 24.0%).

### Sterilising males

Compared to female-only strategies, additionally targeting male FRDs generally had higher population reduction at a similar cost (Fig. 2). Sterilising males affected the population via two routes, reduction of pregnancy rates and catching logistics. With male sterilisation impact (MSI) of 0, 0.15, and 0.25, 70% male sterilisation coverage gave peak pregnancy rates of 10%, 9.0%, and 8.3% respectively, which then had population impacts (Fig. S4). A sensitivity analysis showed that whilst the value of MSI did change population outcomes, this was comparable to or lower than changes in other parameters (Fig. S5). When MSI was 0 (no effect of male sterilisation on pregnancy rates) and teams were reduced based on number of dogs captured, sterilising males still reduced the population more than female-only strategies due to catching logistics. As both male and female FRDs were eligible for capture, catching teams were active for longer periods and more females were sterilised, rather than stopping when less than 5 or 10 females were captured (Fig. S4). When catch returns governed whether sterilisation would continue, the decision to sterilise males was more impactful. For example, in effective strategies based on catch returns, male-inclusive strategies achieved maximum FSC more quickly (median = 212 days, IQR = 151-249 days) than female-only strategies (median = 3601 days, IQR = 3580-3624 days). Whereas in effective strategies based on fixed sterilisation periods female-only and male-inclusive strategies had similar median days to max FSC (F-O median = 2566, IQR = 1826-3551, M-I median = 2872 days, IQR = 1826=3618 days).

### Fixed strategies: Duration, interval, frequency and intensity

None of the continuous sterilisation strategies were optimal. Single sterilisation sessions of varying periods were also never optimal; done for less than 730 days it was ineffective, whereas periods longer than 730 days (excluding independent-only strategies) were effective but high-cost (Fig. 5). Of the fixed-maintain and fixed-reduce strategies (defined number of days between sterilisation periods), the highest proportion of optimal strategies were associated with sterilisation periods lasting 365 days (4.23% optimal strategies), intervals of 365 days (3.85%) or 730 days (3.33%), and 3 or 5 starting teams (3.36% and 3.17% respectively), but these were not statistically significantly different from other values of that parameter. Strategies with sterilisation durations of more than 365 days and intervals of more than 1825 days were less likely to be optimal (Table S3). There were trade-offs between duration, interval, frequency and intensity. For example, in a male-inclusive strategy with five repeats and 1 team, durations of 90 days were ineffective at all intervals tested (90, 185, 365, 730 days). With 3 teams maintained, durations of 90 days were optimal at 185, 365 and 730 day intervals, but not 90 day intervals, and at 5 teams maintained, 90-day durations were effective but high-cost (Fig. S6). Shorter sterilisation durations of 30 days were mainly ineffective (apart from with more than six repeats), whereas longer durations (> 730 days) were high-cost but typically effective (Fig. S6).

**Figure 5.**
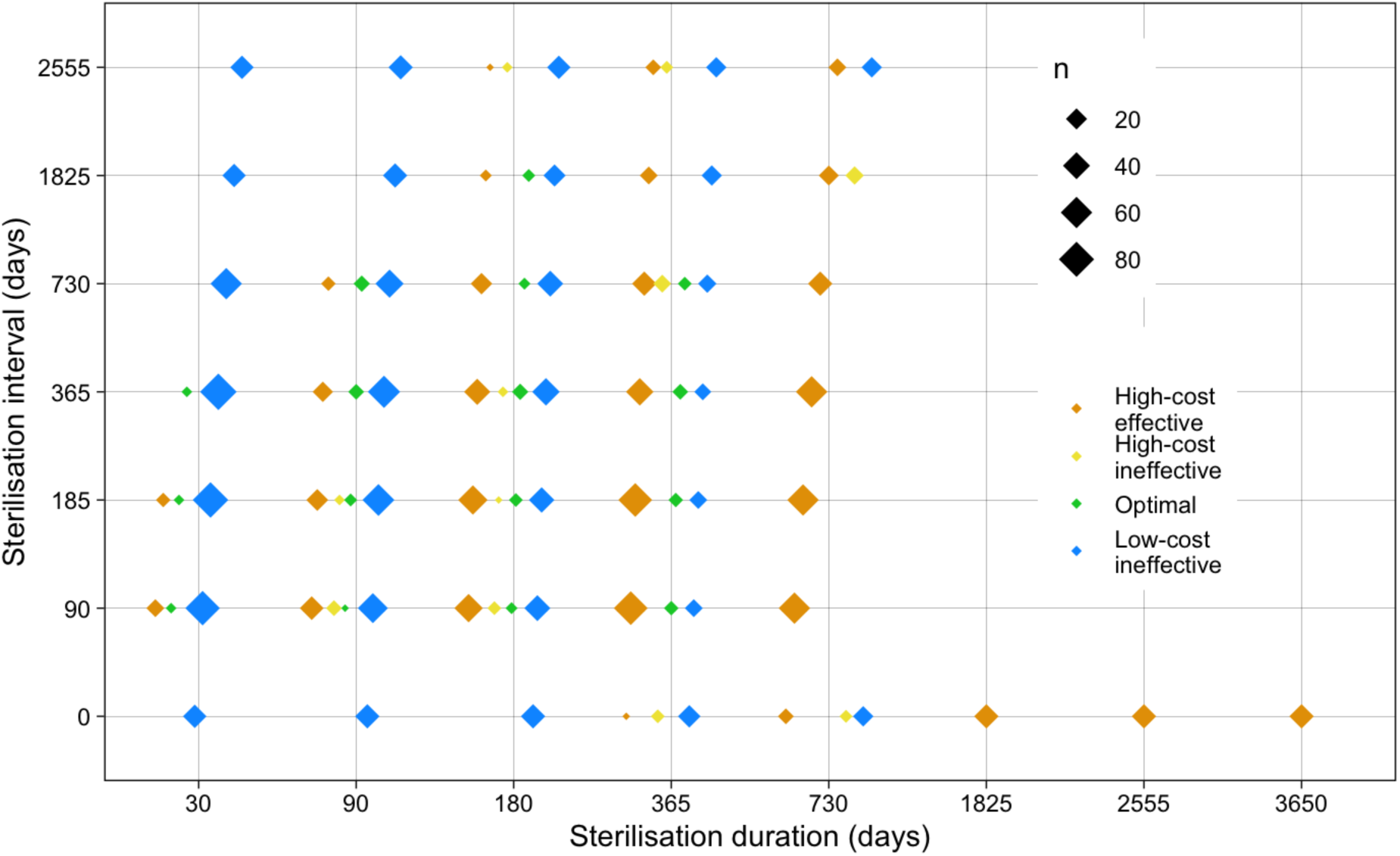
Relationship between sterilisation duration and sterilisation interval, for 600 strategies applied to the free-roaming dog (FRD) population model that sterilised both dependent and independent FRDs. Point size indicates the number of strategies, colour indicates the success of the strategy based on cost (high > US$250k, low <= 250k) and population density reduction (effective = more than 40% density reduction and less than 50% population at 10 years), with low-cost effective strategies being classed as ‘optimal’. Multiple outcomes appear for some combinations due to additional parameters being different, these are explained further in the text and in supplementary figure S6).

Compared to maintaining all teams for the whole simulation, reducing teams based on catch returns meant longer sterilisation durations were more often effective, and no thirty-day sterilisation durations were effective. Though overall there was little difference in the proportion of optimal strategies whether teams were maintained or reduced (3.05 c.f. 2.22%, Table S3), fixed strategies where teams were maintained were more likely to be effective (52.5% c.f. 34.2%). Strategies starting with three or five teams had a higher proportion of optimal strategies than starting with one team (3.36% and 3.17% c.f. 0.91% for one team; Table S3). Starting with more teams achieved a higher sterilisation coverage quicker but with diminishing returns for each additional team and as sterilisation coverage increased (Fig. S7). Starting with more teams increased costs, and this was further demonstrated for longer sterilisation periods, especially where teams were maintained. However, if teams were reduced, this was reduced and having 3 or 5 teams had similar costs.

### Catch-guided strategies: Duration, interval, frequency and intensity

In catch-guided strategies intervals were fixed but sterilisations ceased when dog captures fell below a threshold of 5 or 10 dogs. This resulted in mean sterilisation durations (of multiple sterilisation sessions over the simulation) in optimal strategies between 3–187 days (median = 22, IQR = 13–47 days). Longer intervals gave more time for unsterilised dogs to grow and breed and so produced longer sterilisation durations to ‘catch up’ (Fig. S8) and vice-versa, as longer intervals required fewer repeated strategies. All interval lengths resulted in some optimal strategies. In male-inclusive, dependent-inclusive optimal strategies, a catch threshold of ten produced sterilisation durations of less than 60 days at all intervals, whereas a catch threshold of five produced optimal strategies of durations over 100 days only with longer intervals of 730 or 1825 days (Fig. S8). Female-only optimal strategies with a catch threshold of five had shorter durations (<30 days) and shorter intervals (<=365 days) and there were no optimal female-only strategies with a catch threshold of ten.

As teams were not reduced in catch-guided strategies, a catch return threshold of five effectively meant that when five teams are deployed, catching stopped when each team caught one dog (max FSC = 0.79), whereas when one team was deployed, they stopped when that team caught five dogs (max FSC = 0.55). This model feature resulted in catch-guided strategies with more teams achieving higher sterilisation coverage. Catch-guided strategies with only one team deployed never captured enough dogs to be effective, as sterilisations ceased early on. When selecting only optimal strategies without a rising population at the end of the simulation (population density slope <= 0), thirteen out of eighteen strategies were catch-guided (Fig. S9).

### How many should we sterilise? Number of sterilisation days and proportion of the starting population sterilised

In optimal strategies the total (not consecutive) number of days on which dogs were sterilised was between 270 and 1095 days (median = 450 days, IQR = 327-1046 days). Sterilising on more than 1260 days was exclusively high-cost and sterilising for less than 270 days was ineffective. Over the whole simulation period, optimal strategies sterilised between 102 and 146% of the initial starting population of juvenile and adult females (median = 122%, IQR = 111-133 days). Of the strategies that sterilised more than 150% of the starting population over more than 1000 days, 92% were high-cost ineffective independent-only strategies (Fig. 6).

**Figure 6.**
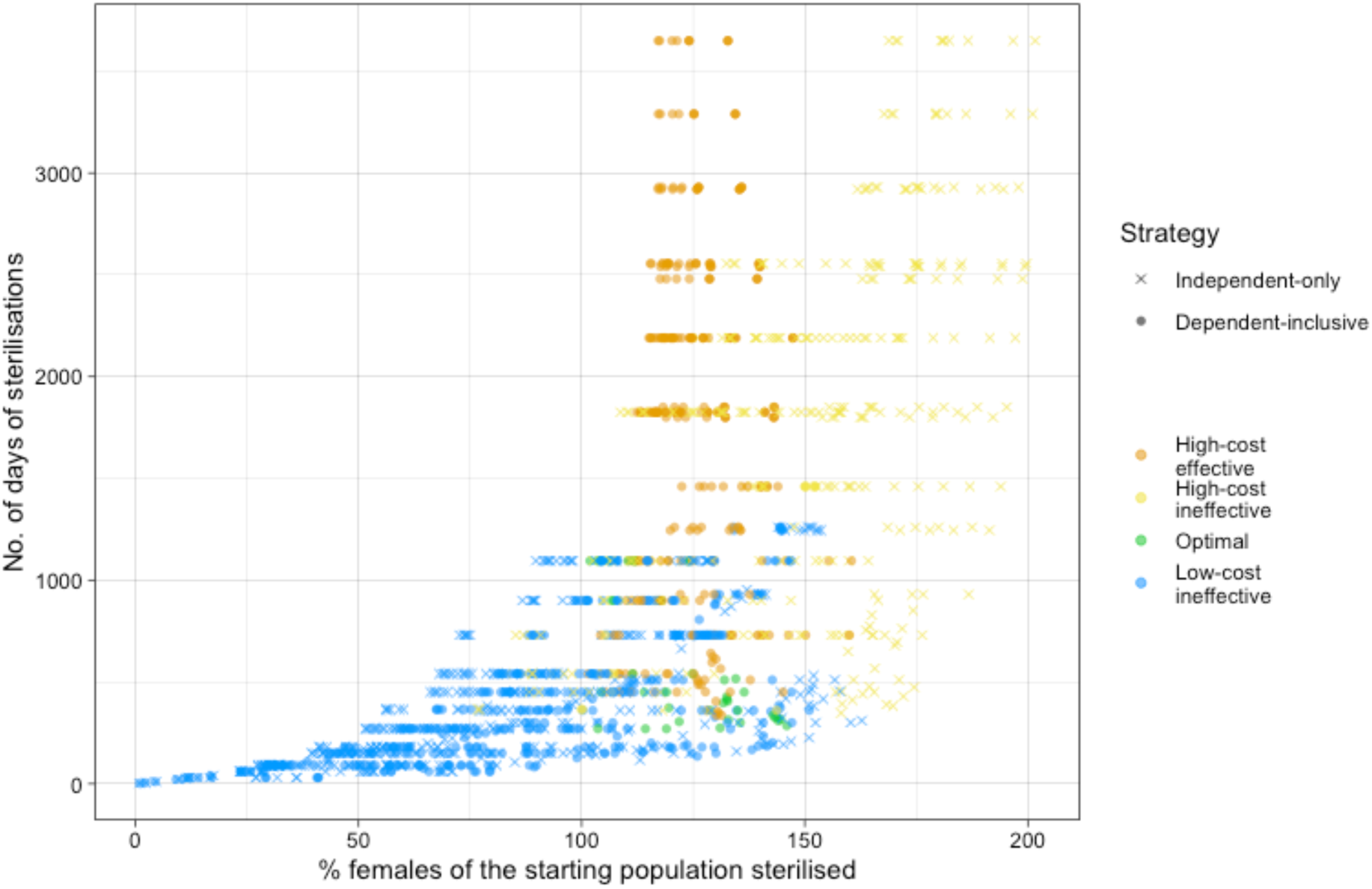
Sterilisation days and total sterilised. Total number of days of sterilisations and percentage of the starting adult and juvenile female population that was sterilised over the 10-year simulation of 1524 sterilisation strategies. Colour indicates the success of the strategy based on cost (high > US$250k, low <= 250k) and population density reduction (effective = more than 40% density reduction and ineffective = less than 50% population at 10 years), where low-cost effective strategies are classed as ‘optimal’. Shape indicates whether the dependent (cared-for by humans) free-roaming population was sterilised.

### What time of year should we sterilise?

Six-month sterilisation sessions with an interval of six months were applied to the model starting in each month of the year to see if there was any effect of timing sterilisations relative to the breeding season. The starting month had minimal influence on the population density decrease (range = 0.417-0.424, mean = 0.421, SD = 0.003) or cost (range = 201.10-202.07k, mean = 201.56, SD = 0.325). However, strategies that start during or shortly after the breeding season increased the number of dogs sterilised during pregnancy or in oestrus (Fig. S10).

### Distribution of costs

Total mean costs were the highest at the start of most strategies (year 1 = $86.8k c.f. year 10 = $13.4k, Fig. S11), reflecting that most sterilisations occurred in year one, with effective strategies having a higher proportion of sterilisations in the first year (effective = 0.67, ineffective = 0.55, *t* = 11.93, *p* = <0.001). Cost per sterilisation was lowest at the start of strategies (year 1 = $48.39, c.f. year 10 = $461.85 per sterilisation). In high-cost strategies in year one, sterilisations cost significantly more per sterilisation in ineffective strategies compared to effective ones (high-cost ineffective (HC-I) = $74.51, high-cost-effective (HC-E) = $58.93, *t* = 4.45, *p* = <0.001) despite year one total costs being similar (HC-I = $136.2k, HC-E = $145.0k, *t* = −1.50, *p* = 0.14). In low-cost strategies in year one, the difference in cost per sterilisation between optimal and ineffective strategies was not significantly different (optimal (O) = $34.42, low-cost-ineffective (LC-I) = $37.37, *t* = 0.99, *p* = 0.33), but year one costs were higher in optimal strategies compared to low-cost ineffective strategies (O = $93.8k, LC-I = $47.9k, *t* = −10.61, *p* = <0.001). On average over the simulation, it would cost $1846 to reduce the population by 100 dogs for 1 year for effective strategies and $2120 for ineffective strategies (cost effectiveness ratio for effective strategies ($ per dogs/day decrease) = 0.051, ineffective = 0.058, *t* = −3.69, *p* = 0.0002). The maximum number of vets was wholly determined by the number of teams deployed, one team required two vets, three teams required four vets and five teams required a maximum of six vets. However, most days of each strategy had only one vet operating.

### Life expectancy, deaths and vaccinations

Life expectancy of independent FRDs increased as FSC increased and as the population decreased below carrying capacity (Fig. S12). In strategies that only sterilised independent dogs, life expectancies were slightly higher for a given population reduction, though at higher population decreases that independent-only strategies could not reach, the life expectancy was higher for male and female independent FRDs (Fig. S12). In independent-only sterilisation strategies, all the sterilisation capacity is directed towards the independent population achieving a higher independent sterilisation coverage and longer life expectancy (Fig. S12). Given that all tested strategies reduced population to some extent, all strategies reduced the number of dogs that would need to be vaccinated in a rabies control campaign (Fig. S13).

### Strategy visualisation – interactive online dashboard

To explore the impacts of individual strategies on sterilisation coverage and population size over time, we created an online interactive dashboard available at https://field.shinyapps.io/DogPopSimApp using R Shiny (Chang et al., 2024). The app allows users to select parameters of strategies tested in this paper and visualise the total, dependent, and independent population size over time alongside the number of sterilisations performed per day, the female and male and dependent and independent sterilisation coverages over the ten-year simulation. Users can select up to three strategies concurrently and can visualise where each strategy lies compared to others in terms of cost and population reduction.

## Discussion

This study tested a wide range of realistic sterilisation strategies, defined by field data, to support decision making within FRD population management strategies. Strategies were evaluated based on their effects on population size and cost. Achieving and maintaining high FSC was strongly associated with effective population reduction and was one of the most influential determinants of success. No single value for sterilisation duration, interval, or frequency defined an effective strategy in isolation; instead, these parameters were interdependent and could be adapted to context, provided they resulted in high mean-daily FSC. Additional male sterilisation enhanced population reduction at little additional cost, due to catching logistics and an assumed relationship between pregnancy rates and male sterilisation coverage. Excluding large demographic groups of females, such as dependent FRDs, constrained sterilisation coverage and limited population reduction. Our model explicitly incorporated seasonal reproduction observed in Indian FRD populations [18] and the timing of interventions relative to this season are likely to have implications for the health and welfare of female dogs, yet had little influence on overall strategy effectiveness. Although the interventions are realistic, when making our conclusions, we acknowledge and account for the important limitation that the population model is closed, and that variations in efficacy are inevitable when applying strategies in real-world settings.

### Sterilisation coverage

Mean-daily FSC and maximum FSC were both good indicators of population decrease, with mean-daily FSC better discriminating optimal strategies. Although mean-daily FSC is difficult to calculate in real time, a rolling mean FSC may provide a practical proxy for monitoring progress. Maximum FSC (63–84%) and mean-daily FSC (47–64%) associated with optimal strategies were mostly in-line with the 70% coverage recommended in India’s Animal Birth Control rules and guidelines [5,6]. Although optimal strategies spent around 66% of the simulation time below 63.3% FSC, the oft-quoted 70% goal seems sensible given that in many open populations, compensatory immigration of unsterilised dogs or emigration of sterilised dogs is likely to dilute effective coverage [11] and greater sterilisation intensity, longer durations, or shorter intervals may be required. Whilst our results represent a minimum input under idealised conditions, the broader principles of optimising coverage and timing are likely to remain robust. Use of physical boundaries or planning to sterilise sequentially in neighbouring areas may limit dog-mediated migration and maximise sterilisation efforts [34].

Ecological parameters were derived from a population with low levels of fertility control. Though increased survival at population sizes below carrying capacity was included in the model, we did not include other compensatory mechanisms such as survival and litter size (in addition to migration). Direct effects of sterilisation on survival may be expected to temporarily reduce population reduction as sterilised dogs stay in the population longer. A population below carrying capacity may increase litter size at birth[35,36], however the foetal numbers during pregnancy used for estimating litter size at birth in this study match very closely litter size at birth for Western pet domestic dogs of similar size [37], suggesting there may be physiological restrictions in domestic dogs. Due to limited data on populations with high sterilisation, we limited our simulation to ten years, comparable with some simulations [38] (Yoak et al., 2023), but shorter than others [39]. We assumed no change in the carrying capacity during this time as the study population had been stable for at least 5 years of monitoring. Many factors can impact on the carrying capacity of a region (e.g. waste management, supplemental feeding) and compensatory ecological mechanisms are likely to drive the population back to that carrying capacity or allow other species to take advantage of resources (e.g. cats, rats). Therefore, different levels of sterilisation may be required depending on carrying capacity changes and the extent of compensatory mechanisms.

### How many and when?

In our ten-year simulation, the proportion of female FRDs to be sterilised for an effective strategy was more than 100% of the initial FRD population, which demonstrates the minimum investment and resources likely to be required for population reduction. Strategies that concentrated a high number of sterilisations (102-146% of starting population) into a low number of sterilisation days were more likely to be optimal when combined with other optimal parameter values, but those days should not be concentrated too close to the start or end of the strategy, as these were associated with high-cost ineffective strategies.

### Sterilising dependent FRDs

In our model population, human-dependent FRDs made up one third of the FRD population and omitting this group limited FSC and population reduction, preventing any strategy being effective. The larger this group, the more impact omitting it will have. In African countries with relatively larger dependent FRD populations, sterilisation efficacy may be more significantly affected by owners or guardians electing not to sterilise their dog[40] or programmes that do not target the dependent population. Frequently, private vet clinics are not affordable and either sterilisation campaigns do not sterilise dependent dogs or guardians are reluctant to take-up free sterilisation due to concerns about having their dog returned or the safety and quality of the surgery[41]. Public-private partnerships for managing free-roaming cats, where owners are given a voucher for free or discounted sterilisation, have worked well in the UK and USA[42,43] and may provide a sustainable solution for this gap. Where owners elect not to sterilise their pets due to concerns about behaviour change or wanting to breed, other options may be required, e.g. confinement. Though we did not include confined dogs in our model, this is another potential source of new dogs which may need targeting with fertility control or restriction, if abandonment from this population is high [39,44].

We assumed that capture costs of dependent and independent FRDs were the same as we did not have sufficient data to show otherwise, but it is possible that capture, transport, housing, monitoring and feeding costs may be different when sterilising dependent FRDs. The ineffectiveness of independent-only strategies would be unchanged, but some high-cost effective strategies including dependent FRDs may become optimal if costs are reduced.

### Inaccessible dogs

The proportion of inaccessible dogs we chose (5% of independent and 10% of dependent dogs) was likely to represent a best-case scenario where other capture techniques in addition to net catching may be used to achieve the higher sterilisation coverage, e.g. darting and sedation, cage-trapping, though we did not account for the increased consumable costs of this type of capture. Even considering this, empirical studies have demonstrated that in some areas the proportion of inaccessible dependent dogs is likely to be at least 10% [45] and uncatchable independent dogs may be around 30% in regions where consistent sterilisation occurred over a long period[16]. Our optimal maximum sterilisation coverage was between 63 and 84%, therefore in areas that can’t achieve a maximum FSC of over 70%, successful population reduction may still be possible, albeit challenging. Ultimately, problems accessing substantial proportions of the free-roaming population for sterilisation will impact on the efficacy of sterilisation strategies.

### Sterilising males

Whether to sterilise male dogs in a dog population management programme is a highly debated issue for stakeholders [17] (Collinson et al., 2021). In our model, female sterilisation clearly led to population reduction, supporting previous findings in modelling studies [23,38,46]. Yet, in our model additionally sterilising male FRDs further reduced population size by limiting birth rates and increasing the number of female and male dogs caught, mechanisms not included in previous modelling. Our assumption of the male effect of sterilisation of 0.15 meant that 70% male sterilisation resulted in a 1% reduction in pregnancy rates and was a conservative value based on limited success of inexperienced males and mate choice exhibited by domestic dogs [25–27]. Even if male sterilisation had no impact on pregnancy rates, the larger pool of unsterilised dogs kept catching teams active and thereby increased sterilisation coverage. It is worth noting this effect may also be achieved if males are captured primarily to vaccinate against rabies rather than to sterilise, as is practiced in some sterilisation projects[47]. Sterilising male FRDs requires less time and fewer consumables and if payments for sterilised dogs are equal for males and females, this may improve cost-effectiveness. Without either of these assumptions, there was no population reduction associated with additionally sterilising males. Some organisations have reported up to 20% of castrations experiencing post-operative scrotal swelling[48]. yet rates of this have been low in the authors’ experience (< 0.5%) [11,49]. Empirical data on the impact of male sterilisation coverage on pregnancy and understanding the realities of catching only female dogs [50] would allow more precise predictions and better assessment of the population value of sterilising male dogs. In addition to population effects, the effect of male sterilisation on behaviour alone may be sufficient to include or exclude it from strategies due to the potential impact on movements, disease transmission, and aggressive behaviour [13,51,52]. This represents an important knowledge gap, given that dog bites and attacks and zoonotic disease are frequent motivations for dog population management [53].

### Duration, interval, frequency and intensity

We found that some continuous strategies over 730 days were effective, as previously reported [39], but they offered poor value for money because teams were paid continuously to catch relatively few dogs. In contrast, repeated shorter sterilisation durations tended to be more efficient: in fixed strategies, optimal durations ranged from 30–365 days while in catch-guided strategies, slightly shorter durations of 3–186 days were often optimal. A wide range of intervals between 90 and 1825 days were in optimal strategies, suggesting that it doesn’t so much matter when sterilisation are repeated, but it does matter that they are repeated, with shorter intervals for shorter sterilisation durations. No single duration or interval consistently guaranteed success, indicating that effectiveness depends on how strategy components interact rather than on any one parameter in isolation.

Catch-guided strategies were particularly effective because sterilisation durations adapted to interval length, ensuring that FSC was maintained. They were also more cost-efficient, as teams (typically 3 or 5) ended capture when catches fell below the threshold, avoiding inefficient use of staff time. In these scenarios, teams could be redeployed to neighbouring areas with lower sterilisation coverage, allowing strategies to run sequentially across multiple areas.

In our model, initially deploying 3 or 5 teams was more likely to be optimal, although in some cases team numbers could be reduced as catch numbers declined. Both catch-cut-off thresholds of 5 and 10 produced optimal strategies, effectively ceasing sterilisation at approximately 1–3 dogs per team.

Unlike some fixed strategies, optimal catch-guided strategies sustained FSC and population reductions by repeating sterilisations until the end of the simulation, suggesting they may be particularly effective for long-term control. Models projecting over longer time horizons may be required to evaluate these outcomes. Catch-guided strategies also provide a real-time operational indicator for when to cease sterilisation, although very short sterilisation periods (e.g. 3 days) are unlikely to be feasible in practice, and organisational capacity and community needs must be considered when selecting appropriate sterilisation periods.

### Costs

Effective strategies achieved greater population reduction per unit cost, indicating that optimising impact can also improve cost-effectiveness. In low-cost strategies, higher first-year expenditure was associated with effectiveness, and intensive sterilisation early in the programme was a consistent feature of successful strategies, supporting the value of ‘front-loading’ interventions [54]. Although cost per sterilisation was lowest at the start due to high capture efficiency, total costs were highest because there were many dogs to sterilise. Over time, cost per sterilisation increased as capture efficiency declined; however, in effective strategies this was offset by a smaller pool of unsterilised dogs that were sterilised frequently enough to sustain population reduction. Optimal catch-guided strategies with intervals of less than 365 days had better cost effectiveness ratios and lower overall costs than other strategies, suggesting that annual or more frequent sterilisation periods may be recommended. Ineffective low-cost strategies failed due to insufficient sterilisation, whereas ineffective high-cost strategies sterilised more dogs but achieved limited population impact because poorly timed interventions (e.g. short durations or long intervals) allowed population recovery.

### Seasonality

Although reproductive seasonality has been identified in FRD populations, it has rarely been incorporated into population models. Our model explicitly accounted for this pattern and tested starting strategies in each month of the year. Though it had limited effect on population reduction, some start months increased the number of females sterilised during high-risk periods of oestrus and pregnancy. This is likely to result in negative effects not included in the model such as a detrimental impact on animal welfare, increased mortality of unborn pups, and increased surgical or post-operative recovery time due to highly-vascularised tissues. Timing of sterilisation should therefore prioritise practical and animal welfare factors, e.g. avoiding very wet seasons and peak breeding and pregnancy periods.

### Strategy planning

The Animal Welfare Board of India advise that catching teams ‘work systematically to sterilize at least 70% of the dog population in a time bound and area based manner’ [5], this was practiced by our teams in the field and is represented in our model. To achieve this, we mapped boundaries of the intervention area and teams were guided by these boundaries, similar to the approach used in rabies vaccination [55], i.e. teams were instructed to return to areas with low coverage. Whilst this is essential for achieving higher coverage, it can be frustrating and unrewarding for catching teams to keep returning to areas with only harder-to-catch dogs and incentivising high coverage rather than dog numbers may be useful here. Within a designated area, population size and sterilisation coverage can be assessed to estimate costs of an effective strategy. This is likely to be an iterative process where available costs are balanced with the desired area and magnitude of impact, and a compromise is reached. The model was grounded in the reality of catching FRDs in an urban area of Goa and simulated an approach where veterinary teams were solely focussed on sterilisation, as this is the most common experience of the authors. We suggest that the initial high-volume of sterilisation is completed by such teams (e.g. five catching teams and up to a maximum of six vets in a similar-sized area and dog density) but other options for maintaining the initial effort may be to utilise existing local veterinary capacity (by upskilling government veterinary dispensaries or public-private partnerships) [41] combined with roving catching teams, thus reducing the overheads and veterinary costs when dog captures are low and only one vet is required, and reducing the high cost per sterilisation towards the end of strategies. Though absolute numbers are only applicable in our specific ecological context, users of our online dashboard can better understand the population impacts of different approaches with an aim to facilitate evidence-based decision making. Allowing manipulation of the simulated population will be the focus of future work. Realistic, data-driven planning of campaigns manages expectations and allows limited resources to be directed towards more-effective strategies.

### Wider benefits and limitations of sterilisation

Contribution from increased survival of FRDs at population sizes lower than carrying capacity (intrinsically included in the model) enhanced life expectancy and reduced deaths. Whilst not included in the model, sterilised FRDs are also reported to have better condition, longer survival and fewer infectious diseases than their unsterilised counterparts [11,12,22,56]. Decreased barking in areas with higher levels of sterilisation has been reported, which may have positive benefits for the local community [11].

Historically, reduced population density by culling has not been successful in controlling rabies[57,58], yet reduced population turnover and increased longevity, achieved with sterilisation in our model, is likely to maintain rabies immunity for longer and reduce vaccination requirements, as was found in a rabies modelling study based on data from Tamil Nadu [7]. Rabies vaccination is routinely given to dogs captured for sterilisation in India, therefore some strategies are likely to save additional costs by removing the need for a separate anti-rabies vaccination programme, if they maintain immunity over the critical vaccination threshold for rabies [59]. Sterilisation may increase immigration where sterilisation areas are small [11], however this is likely to be on a much smaller scale than the perturbation effect seen in culling [60,61] and can be ameliorated with larger or geographically contiguous sterilisation areas [34]. Surgical sterilisation provides permanent and 100% effective fertility control, but is costly and labour intensive. Injectable contraceptives may be suitable in the future which would dramatically increase the feasibility of fertility control in FRD populations[62] and reduce risk of dog-dog disease transmission and potential stress of transport to sterilisation centres [63].

Despite our focus on sterilisation in this study, we support the view that sterilisation needs to be combined with other interventions to create a holistic dog population management strategy as recommended by WOAH[4]. Concurrent implementation of sterilisation and ‘responsible dog ownership’ (decreased abandonment and increased adoption from shelters) proved to be more effective at reducing the FRD population than the sum of both individually, particularly at 70% sterilisation coverage, therefore this is likely to be a powerful, cost-effective and animal-welfare-positive combination, though this assumed a 90% change in abandonment and adoption [39]. However, we should be cautious in using the term ‘responsible dog ownership’ interchangeably between cultures, our research shows that dogs adopted from the street in India are usually still allowed to roam [11] and may not actually decrease the FRD population. In addition, sheltering or housing dogs accustomed to free-roaming may compromise their long-term welfare [64]. The term ‘ownership’ may also be problematic for free-roaming dogs because it implies an individual responsibility, whereas in many settings human-dog relationships may be better characterised by care, guardianship or community-level responsibility. Measures that combine population reduction and reducing carrying capacity, e.g. limiting food/waste resources are also likely to be particularly sustainable, however evidence is lacking [65].

## Conclusion

Sterilisation can form part of a sustainable approach to dog population management, and stakeholders planning and implementing this intervention can optimise strategies to maximise the impact of each rupee or dollar spent. Low levels of sterilisation reduced population turnover and puppy mortality and may benefit rabies immunity and individual animal welfare; however, for population reduction, sterilising for short durations with no or very infrequent follow-up is likely to be ineffective, costly, or both, especially where compensatory mechanisms occur, e.g. migration. Short sterilisation durations repeated at appropriate intervals to maintain moderate FSC were optimal for population reduction and cost-efficiency. Targeting dependent FRDs was essential to population reduction, particularly where they constitute a large proportion of the population. Additionally sterilising male FRDs is likely to be cost-effective, assuming an effect on pregnancy rates. These findings highlight the need for planned, coverage-driven sterilisation strategies that prioritise early intensity and appropriate targeting, rather than maximising the number of sterilisations delivered or adhering to fixed schedules.

## Data availability statement

R code used for the simulation model, model processing, visualisation, and Shiny web application is publicly available at: https://github.com/fieldingh/DogPopSim. A permanent archived version with DOI will be provided upon acceptance.

## Acknowledgements

HRF was supported by a Dogs Trust Canine Welfare Grant (ID R45810) and Worldwide Veterinary Service (WVS). SM is funded the Biotechnology & Biological Sciences Research Council Institute Strategic Programme grant funding to the Roslin Institute (BBS/E/RL/230002D). The funders played no role in the design of the study and collection, analysis, and interpretation of data and in the writing of the manuscript. This work has made use of the resources provided by the Edinburgh Compute and Data Facility (ECDF) (http://www.ecdf.ed.ac.uk/). For the purpose of open access, the author has applied a CC-BY public copyright license to any Author Accepted Manuscript version arising from this submission.

## Author Contributions

HRF: Conceptualization, Data curation, Formal analysis, Investigation, Project administration, Methodology, Visualisation, Writing – original draft, Writing – review & editing. PB: Conceptualization, Methodology, Supervision, Writing – review & editing. AG: Conceptualization, Project administration, Funding acquisition, Methodology, Data curation, Writing – review & editing. KF: Funding acquisition, Methodology, Data curation, Writing – review & editing. AVR: Methodology, Data curation, Writing – review & editing. LG: Funding acquisition, Project administration. BMCB: Funding acquisition, Supervision, Writing – review & editing. IH: Funding acquisition. RK: Funding acquisition, Writing – review & editing. RJM: Conceptualization, Funding acquisition, Project administration, Supervision, Writing – review & editing. SM: Conceptualization, Methodology, Funding acquisition, Supervision, Writing – review & editing. All authors have read and agreed to the final version of the manuscript.

